# High-resolution Cryo-EM visualises hydrated type I and IV pilus structures from enterotoxigenic *Escherichia coli*

**DOI:** 10.1101/2024.11.14.623551

**Authors:** Kazuki Kawahara, Hiroya Oki, Minato Iimori, Ryuki Muramoto, Tomoya Imai, Christoph Gerle, Hideki Shigematsu, Shigeaki Matsuda, Tetsuya Iida, Shota Nakamura

## Abstract

To efficiently colonise intestinal epithelium, enteric pathogens utilise a wide variety of pilus filaments, including those synthesised by the chaperone–usher or type IV pilus assembly pathway. Despite the importance of these filaments as potential drug and vaccine targets, their large size and dynamic nature present challenges for atomic-resolution structural determination. Here, we used cryo-electron microscopy (cryo-EM) and whole genome sequencing to identify and determine the structures of type I and IV pili expressed in enterotoxigenic *Escherichia coli*. The well-defined cryo-EM maps at resolutions of 2.20 and 1.78 Å for Type I and Type IV pilus, respectively, facilitated *de novo* structural modelling for these filaments, revealing side chain structures in detail. Further, we resolved thousands of hydrated water molecules around and within the inner core of the filaments, functioning as a buffer to stabilise the otherwise metastable subunit assembly. The high-resolution structures offer novel insights into the subunit–subunit interactions, providing important clues to understand pilus assembly, stability, and flexibility.

## Introduction

Enteric bacterial pathogens have evolved numerous surface organelles to efficiently colonise their host environment (Edwards and Puente, 1998; Kline et al., 2009). Most of these organelles are filamentous protein polymers termed pili or fimbriae, composed of thousands of subunits called pilins (Hospenthal et al., 2017). For enterotoxigenic *Escherichia coli* (ETEC), a major cause of diarrhoea in travellers and children in developing countries, a variety of pilus filaments have been identified, most of which are colonisation factors (CFs) (Madhavan and Sakellaris, 2015). The CFs are categorised into either CF antigens (CFAs) or coli surface antigens and classified into two pilus types synthesised by distinct assembly mechanisms, chaperone–usher pilus (CUP) and type IV pilus (T4P) assembly pathways (Craig et al., 2004; Craig et al., 2019; Giltner et al., 2012; Hospenthal et al., 2017; Madhavan and Sakellaris, 2015; Pelicic, 2023; Waksman and Hultgren, 2009). In the CUP assembly pathway, major and minor pilins are polymerised via the extensively studied “donor-strand exchange” mechanism, in which an incomplete immunoglobulin (Ig)-like fold of each pilin subunit is complemented by a donor strand provided by the N-terminal extension of an adjacent subunit (Hospenthal et al., 2017; Madhavan and Sakellaris, 2015; Waksman and Hultgren, 2009). Meanwhile, the T4P assembly pathway is substantially more complex than the CUP assembly pathway (Craig et al., 2004; Craig et al., 2019; Giltner et al., 2012; Pelicic, 2023). It involves spanning supramolecular machinery, consisting of multiple membrane proteins that function concertedly to assemble each pilin subunit, utilising mechanisms that have yet to be fully understood.

As the CUP and T4P play crucial roles in bacterial infection, they are key targets in the post-antibiotic era for developing novel vaccines and antiadhesive drugs against multi-drug resistance pathogens (Rasko and Sperandio, 2010; Steadman et al., 2014; Wizemann et al., 1999). However, the large size and insoluble and non-crystallising nature of these filaments pose challenges for the implementation of conventional high-resolution structural biology methods such as X-ray crystallography and nuclear magnetic resonance (Egelman, 2017; Garnetta and Atherton, 2022). Therefore, structural characterisation has been primarily confined to the pilus subunit level using either detergent-solubilised full-length pilins (Craig et al., 2004; Craig et al., 2019; Giltner et al., 2012; Pelicic, 2023) or modified pilin constructs to improve protein solubility (Waksman and Hultgren, 2009). The filament structure was then constructed by docking these high-resolution monomeric structures into medium to low-resolution (4.0–15.0 Å) density maps obtained by negative-stain or cryo-electron microscopy (cryo-EM) structural analyses (Bardiaux et al., 2019; Craig et al., 2006; Hospenthal et al., 2017; Kolappan et al., 2016; Li et al., 2012; Spaulding et al., 2018; Wang et al., 2017).

Owing to the technological advancements of cryo-EM, the resolution has recently been much improved (Garnett and Atherton, 2022). Near atomic resolution (∼3.0 Å) structures, even for bacterial filaments such as CUP and T4P, are beginning to be reported, providing new insights into the intact pilus structures for both pilus types (Doran et al., 2023; Neuhaus et al., 2020; Sonani et al., 2023; Treuner-Lange et al., 2024). Nevertheless, the dynamic and flexible natures of these filaments still limit the achievable resolution, and the EM density maps obtained in previous studies are insufficient to define the exact positions of amino acid side chains or to locate solvent molecules. Thus, key biophysical features such as pilus assembly, stability, and flexibility remain to be elucidated.

Here, we determined the cryo-EM structures of CUP and T4P from wild-type ETEC strain 31-10 at resolutions of 2.20 and 1.78 Å, respectively, the highest-resolution structures reported to date for each pilus filament family. The quality of density maps enabled us to perform *de novo* structural determination of these filaments and reveal unprecedented atomistic details of subunit– subunit interactions involving thousands of hydrated water molecules resolved for the first time during bacterial filament structural analysis. The high-resolution cryo-EM structures provide insights that enhance our understanding of the biology of CUP and T4P filaments, both of which play crucial roles in bacterial colonisation and pathogenesis.

## Results

### Purification of the filaments from ETEC 31-10

ETEC strain 31-10 cells were grown on a CFA agar plate for optimal expression of CFs (Evans et al., 1977). We used culture conditions previously shown to be well suited for the efficient expression of CFA/III, one of the CFs belonging to a member of the type IVb subclass of T4P (T4bP), which is responsible for the ETEC adherence to intestinal epithelium (Kawahara et al., 2016; Oki et al., 2018). Transmission electron microscopy (TEM) analysis of the purified filaments based on negative staining reveals one dominant filament type with a diameter of approximately 80 Å and highly flexible morphology (Appendix Figure. S1), consistent with the characteristics observed for the CFA/III filament (Kawahara et al., 2016; Kolappan et al., 2012; Taniguchi et al., 2001).

### Cryo-EM of the type IVb pilus from ETEC strain 31-10

In contrast to its negative-staining TEM images, the cryo-EM micrographs showed that the filament has an unexpectedly straight morphology (Fig. 1a). The collected cryo-EM movies were processed and analysed using CryoSPARC (Appendix Figure. S2) (Punjani et al., 2017). The 2D class averages of the filament segments showed a homogeneous class amenable to performing reconstruction at a resolution of 2.52 Å following the application of an asymmetric helical refinement (Fig. 1a). The symmetric parameters were then estimated, and one clear solution, with a rise of 8.01 Å and a twist of 94.63° was found. Subsequent helical reconstruction using these parameters followed by several cycles of contrast transfer function (CTF) and helical refinements resulted in a high-quality cryo-EM density map with 1.78 Å resolution (Fig. 1b). This surprisingly high-resolution reconstruction differs markedly from the previous cryo-EM reconstructions of T4Ps, in which the substantial filament polymorphism severely limits the achievable resolution (Garnett and Atherton, 2022).

**Fig. 1.**
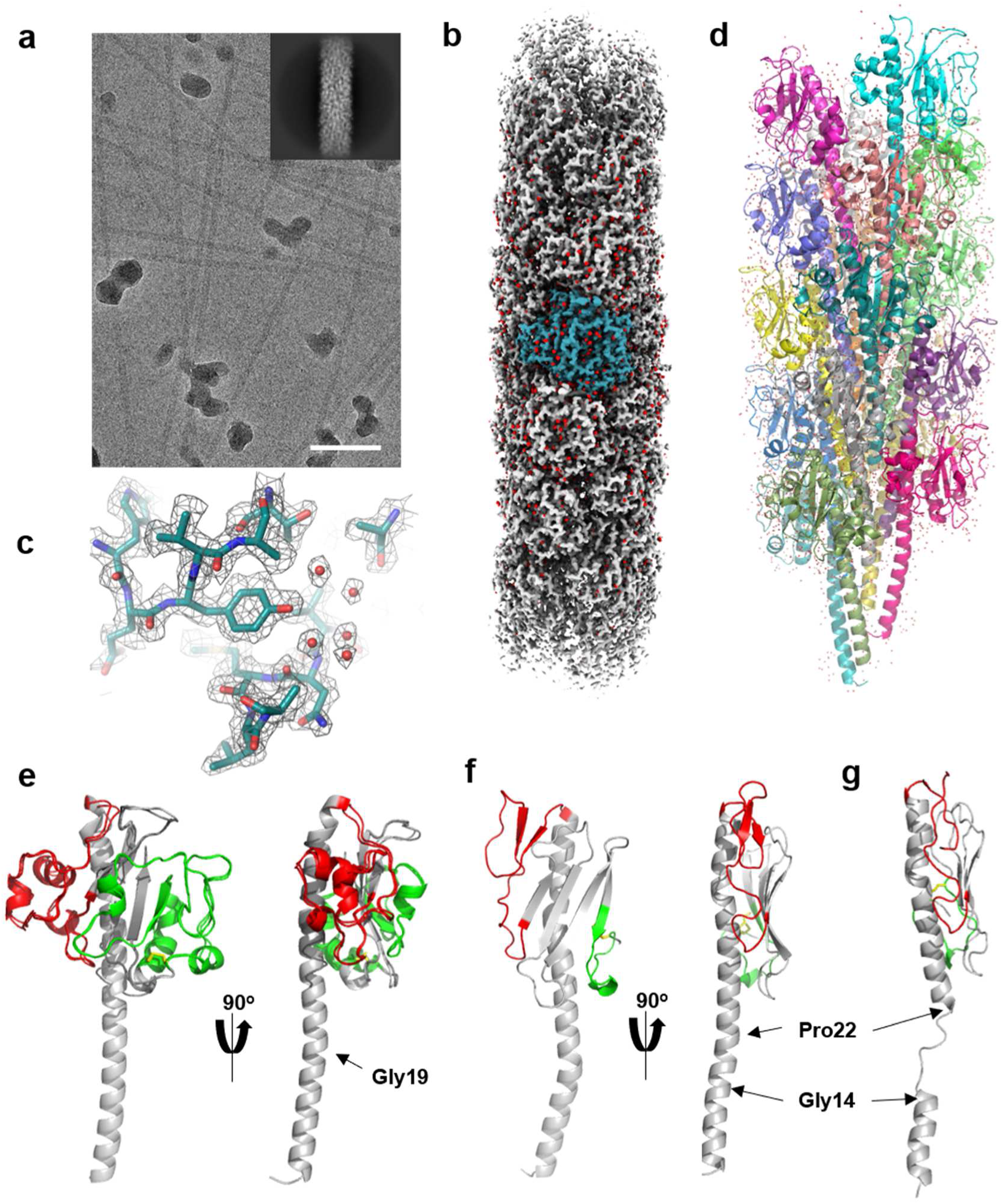
Cryo-EM structure of CFA/III from ETEC strain 31-10. **a**, A representative example of a cryo-EM micrograph showing a highly straight filament with a diameter of 70–80 Å. A representative image of 2D class averages is shown in the inset. The scale bar represents 50 nm. **b**, The cryo-EM density map of the CofA filament. The densities corresponding to water molecules are coloured in red. A part of the density map corresponding to one CofA subunit is shown in deep teal. **c**, A close-up view of the EM density map around Tyr145. **d**, Ribbon representation of the CofA filament model composed of 21 pilin subunits. **e**, Superposition of full-length structure of the CofA subunit in the filament (grey) and crystal structure of ΔN28-CofA (PDB code: 3VOR) (dark grey). The αβ-loop and D-region are coloured in red and green, respectively. In CofA, the αβ loop connecting α1 and the central β-sheet consists of residues 54 to 104, and the D-region located at the C-terminus and stabilised by a Cys132-Cys196 disulfide bond consists of residues 132 to 196, surrounding the αβ core structure. The position of conserved Gly19 is indicated by a black arrow. The disulfide bond Cys132-Cys196 in the D-region is shown as a yellow stick model. **f, g,** Full-length structure of the major pilin PAK from *Pseudomonas aeruginosa* in crystal (**f**) or in filament (**g**) form (PDB code: 1OQW and 5VXY, respectively) as a representative example of T4a major pilin. The αβ-loop and D-region are coloured in red and green, respectively. The positions of Gly14 and Pro22 are indicated by arrows. The disulfide bond Cys129-Cys142 in the D-region is shown as a yellow stick model.

The density map exhibited sufficient quality to construct a filament model *de novo* using the program ModelAngelo without requiring any sequence input (Fig. 1c) (Jamali et al., 2024). The estimated subunit sequence derived from the initial model was ∼88% matched to the sequence of a major pilin CofA from CFA/III (GenBank code: BAB62897.1) (Taniguchi et al., 2001). Subsequent use of the sequence results in a full-length CofA model (208 residues) for each pilin subunit in the filament (Fig. 1d, e). Strikingly, the resolution of the reconstructions enabled in locating 4,175 water molecules distributed both at the inner and outer surface of the filament (Fig. 1b, c). No residual densities corresponding to post-translational modifications were observed, and the density maps and images of 2D class average did not show any evidence of incorporation of the CFA/III minor pilin, CofB, within the filament body. The latter observation is consistent with the notion that the minor pilin only localises at the pilus tip and may transiently interact with the base of a growing filament to induce pilus retraction (Oki et al., 2022).

Each CofA subunit in the filament adopts a typical αβ-roll pilin fold consisting of one ∼50 residue long α-helix (α1) embedded by a central five-stranded antiparallel β-sheet (Fig. 1d). CofA is one of the largest pilins reported to date and belongs to the type IVb (T4b) subclass (Roux et al., 2012). T4b pilin shares a similar αβ-roll fold with pilins of the type IVa (T4a) subclass but is considerably different from those in the type IVc subclass, which only has an α1 consisting of 40–60 residues as a main structural motif (Denise et al., 2019; Oki et al., 2022). The globular domain of T4b pilin (∼180 to 200 residues) is generally larger than T4a pilin (∼150 to 175 residues), which is attributed to its considerably long variable regions, i.e., αβ-loop and D-region (Giltner et al., 2012) (Fig. 1e, f).

The globular domain of CofA is well superposed onto the crystal structure of the N-terminal 28-residues truncated CofA (ΔN28-CofA) (Cα root-mean-square deviation (RMSD) of ∼0.7Å) (Fukakusa et al., 2012), implying no noticeable conformational change required for filament assembly (Fig. 1e). The structure also reveals that α1 forms a straight continuous helix similar to monomeric X-ray crystal structures of full-length T4aP major pilins (Craig et al., 2003; Craig et al., 2006; Hartung et al., 2011; Parge et al., 1995), but drastically different from the T4aP major pilin structures in the filament, where α1 is partially melted at the conserved helix breaking residues, including Gly14 and Pro22 (Fig. 1g) (Bardiaux et al., 2019; Garnett and Atherton, 2022; Kolappan et al., 2016; Neuhaus et al., 2020; Treuner-Lange et al., 2024; Wang et al., 2017). Although Pro22 is not conserved among T4b pilins (Sonani et al., 2023), three glycines conserved at positions 11, 14, and 19, may contribute to α1 flexibility. Noteworthy, the α1 of CofA is slightly bent at Gly19, which apparently optimises the hydrophobic interactions among α1s at the filament core (Fig. 1d, e).

In the filament, each subunit (S) has contacts with a total of six subunits, S/S_±1_, S/S_±3_, and S_±4_, each along with a right-handed 1-start, a left-handed 3-start, and a right-handed 4-start helix, respectively, as observed in other T4P (Fig. 2a). The filament is primarily stabilised by a helically arranged interactions of α1s, in which the hydrophobic N-terminal half of α1 (α1-N: residues 1 to 25) of each subunit (S) has intimate hydrophobic contacts with the α1-Ns of adjacent subunits (S_±1_) and is also interdigitated by the middle part of the α1s of subunits (S_–3_ and S_–4_) (Fig. 2b,c). The tight hydrophobic interaction along the 1-start helix is a conserved feature of the T4P family filament and is critical for T4P stability by forming a hydrophobic filament core.

**Fig. 2.**
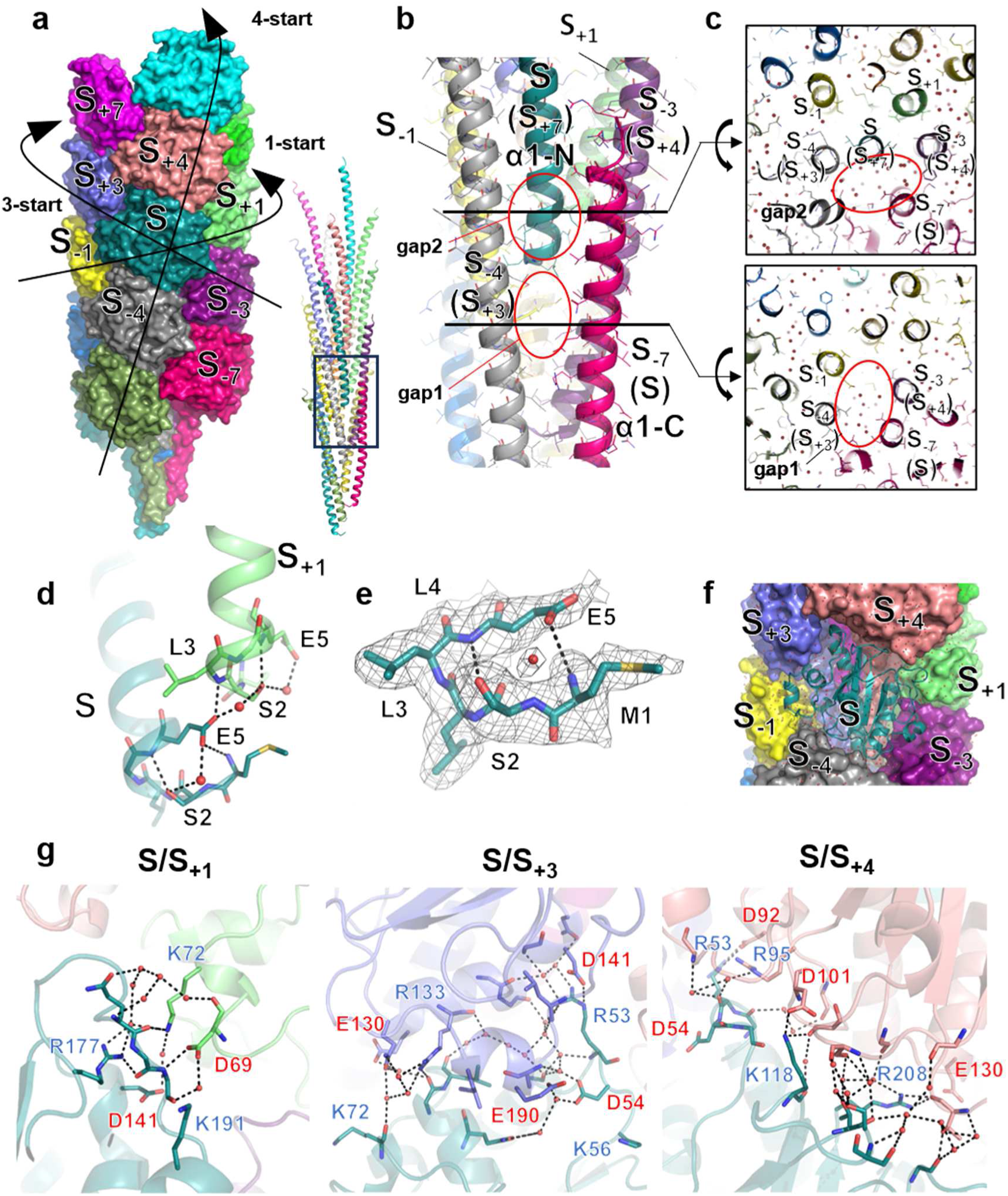
Subunit–subunit interactions in the CofA filament. **a**, Surface representation of CFA/III filament, where each subunit is coloured differently (left panel). One of the subunits coloured in deep teal is a reference and labelled as S. The remaining subunits are labelled along the 1-start helix. The directions of the subunits along the 1-, 3-, and 4-start helices are indicated by arrows. The ribbon representation of α1 interactions is also shown in the right panel. **b,** Close-up view of the α1 interactions. The position of each subunit is labelled, where one of the subunits coloured in deep teal is taken as a reference (S). The labelling is also made by one of the subunits coloured in magenta as a reference (S) and indicated in parenthesis. Two gap regions are indicated by red circles. **c,** Two representative slices of the CofA filament at positions indicated in b, showing pools of water molecules at gap 1 (bottom panel) and gap 2 (top panel) region. **d,** Close-up view of the N-terminus interaction between subunit S and S_+1_ along the 1-start helix. **e,** EM density map of the N-terminal five residues of a subunit at S position contoured at the 2.0 RMSD level. The N-terminal residues are shown as stick models and labelled. A water molecule located between Glu5 and Ser2 is shown as a red sphere. **f,** Molecular packing around subunit S. Each CofA subunit (S) has contacts with a total of six CofA subunits (S_±1_, S_±3_, and S_±4_). **g,** Interactions observed at the molecular interface between each pair of CofA subunits, S/S_+1_, S/S_+3_, and S/S_+4_. For each CofA subunit, the residues involved in the interactions are labelled and shown as stick models. The water molecules involved in the interaction networks are shown as red spheres.

Surprisingly, the intermolecular salt bridge between conserved Glu5 and the N-terminal amine of the adjacent subunit, which is proposed to be crucial in pilus assembly and stability (Craig et al., 2019; Kolappan et al., 2016), is not observed. Instead, the side chain of Glu5 interacts with the backbone amide of Leu3 on the adjacent subunit S_+1_ (Fig. 2d). The density map demonstrated that the N-terminal five residues extend outward from the helical axis of α1 and adopt a turn-like conformation, in which Glu5 intramolecularly interacts with the hydroxyl of Ser2 and the N-terminal amine of Met1 (Fig. 2e). The Glu5 also aids water-mediated interactions with two nearby Ser2 residues, and this relayed interaction further strengthens the α1-N interactions along the 1-start helix (Fig. 2d). We found a water-filled gap (gap1) at the bottom of each α1-N (Fig. 2c). The water in gap1 form hydrogen-bonding networks that run helically through the inner groove of the otherwise hydrophobic filament core (Figs. 2c and 3a). As observed in a recent cryo-EM analysis of the actin filament (Reynolds et al., 2022), the existence of a water channel in the filament not only helps to stabilise the hydrophilic interactions in the filament core but also may lubricate the mechanical rearrangements of subunits upon responding to external forces.

**Fig. 3.**
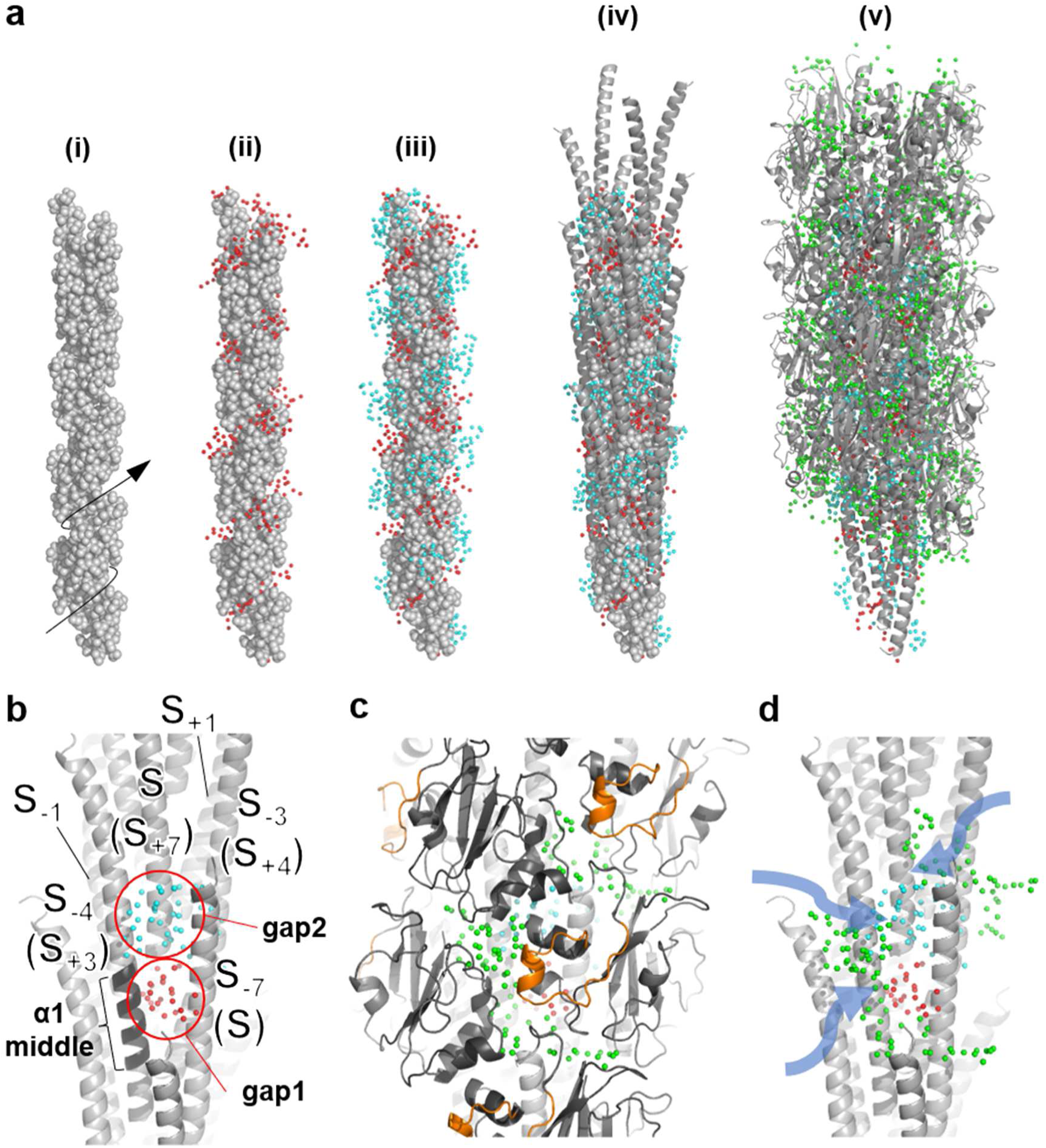
Hydration structure of the CofA filament. **a**, Layers of water molecules in the CofA filament. The sphere representation of the hydrophobic filament core, formed by the interactions among each hydrophobic α1-N (6–16 residues) of the CofA subunit, shows a helically arranged inner groove (i). The inner groove shown by the arrow in the left panel is filled with water molecules (red-coloured spheres) located in the gap 1 region (ii). The gap2 water molecules (blue-coloured spheres) helically cover the outer surface of the hydrophobic filament core (iii). The rest of the α1 (17–51 residues) and globular domains wrapped around the filament core, and the molecular interface of globular domains are filled with a substantial number of interfacial water molecules (green-coloured spheres) (iv and v). **b,** An example of water distribution at two gap regions. Water molecules at gaps 1 and 2 are coloured in red and blue, respectively, and depicted as spheres. A middle region of α1 of one of the CofA subunits (S_-4_ or alternatively say S_+3_ in this case) next to the gap water molecules, is coloured in black. **c,** An example of interfacial water molecules (green spheres) distributed around one subunit (S_–7_ or alternatively say S in this case). The characteristic 3/10 insert (59–78 residues) found in CofA, which partly fills the molecular interface is coloured in orange. **d,** Possible access routes for bulk water to reach into the filament core via communication with water molecules at two gap regions.

The amphipathic C-terminal half of α1 (α1-C: residues 25 to 51) of each subunit (S) is, on the other hand, surrounded by α1s of subunits S_+3_, S_+4_, and S_+7_ (Fig. 2b, c). Although the α1-C (S) forms relatively loose contacts with the α1 of subunit S_+4_, there is a notable gap (gap2) between α1-C (S) and the other two α1s (S_+3_ and S_+7_). Gap2 is filled with water molecules that can communicate with water molecules in gap1 (Fig. 3a,b). The gap water molecules are likely to function to buffer otherwise unstable α1 packing at this subpart of the filament core. Notably, these water-filled gaps are present right next to the middle part of α1 (at S_–4_ and alternatively at the S_+3_ position in Figs. 2b and 3b) and are capable of accommodating conformational changes of α1 (e.g., melting), when it occurs, as observed in other T4P filaments (Bardiaux et al., 2019; Garnett and Atherton, 2022; Kolappan et al., 2016; Neuhaus et al., 2020; Treuner-Lange et al., 2024; Wang et al., 2017).

Owing to the large molecular size of CofA, the filament surface is densely packed with each CofA globular domain (Fig. 2a,f). However, only a few direct contacts involving a limited number of hydrogen bonds, salt bridges, and hydrophobic interactions are formed at each interacting interface (S/S_±1_ or S/S_±3_ or S/S_±4_) (Fig. 2g). The interfaces are instead stabilised by an extensive number of water-mediated hydrogen-bonding networks. The networking water molecules are distributed throughout the filament, enabling bulk water to easily permeate into the filament core, especially via communication with water molecules in the two gap regions (Fig. 3b,c,d). Water accessibility is apparently affected by the packing of globular domains, dictated by the interactions between variable regions. In CofA, a long αβ-loop containing a characteristic 3/10-helix partly fills the interface between subunits, thus making it narrower than the other T4P filament, with relatively short variable regions that would intriguingly expose α1 (Fig. 3c)^18,41^. In addition to the surface-localised hydrated water molecules, these interfacial water molecules could play an important role in pilus stability and flexibility.

### Cryo-EM of the type I pilus from ETEC strain 31-10

In addition to CFA/III, we recognised another straight filament in the cryo-EM micrographs, which has a different morphology from T4P (Fig. 4a). To characterise this filament, we segmented the corresponding filament images and performed iterative rounds of segment picking and 2D classification (Appendix Figure. S3). The 2D class averages showed some features suggestive of CUP (Fig. 4a). We used several symmetrical parameters from previously reported structures of CUP as initial symmetric estimates and found a possible combination of an axial rise of 7.72 Å and a twist angle of 114.90°. Subsequent helical refinement using this parameter set leads to a well-defined density map at a resolution of 2.20 Å (Fig. 4b). The model was created *de novo* using the program ModelAngelo without any sequence input (Punjani et al., 2017), resulting in a filament model, in which each subunit consists of 158 amino acid residues. The BLASTp search using the estimated sequence showed an approximate 84% match to the sequences of type I fimbrial subunit protein FimA from *Shigella flexneri* and ETEC (GenBank code: UMV06456.1 and QED7455.1, respectively), suggesting that the observed filament is a type I pilus (T1P).

**Fig. 4.**
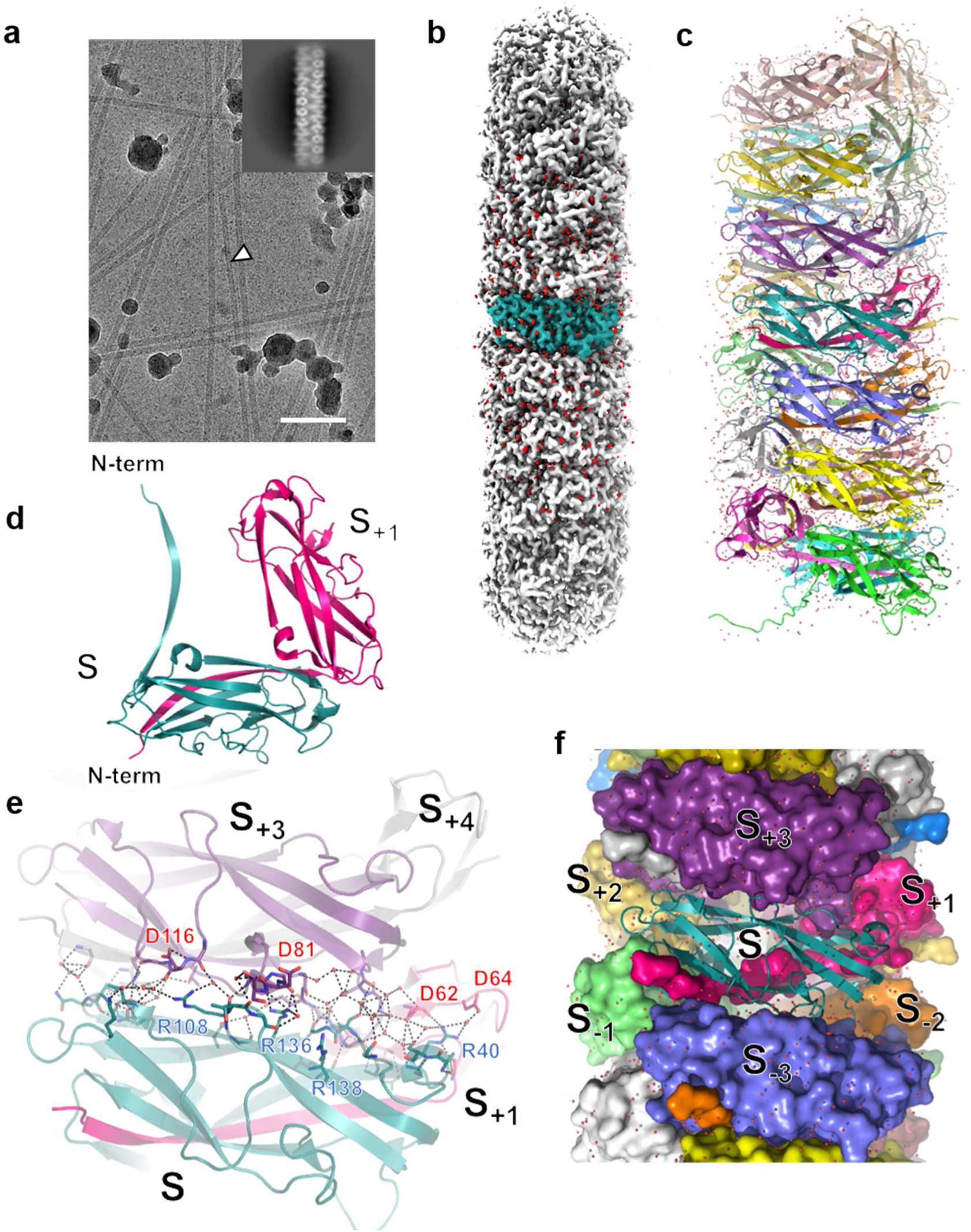
Cryo-EM structure of the type I pilus from ETEC strain 31-10. **a**, An example of cryo-EM micrograph showing the existence of a relatively thin (∼70 Å) filament, which is morphologically different from CFA/III. The filament is indicated by a white arrowhead in the micrograph. A representative image of 2D class averages of the filament showing a feature of T1P is shown in the inset. The scale bar represents 50 nm. **b**, The cryo-EM density map of the FimA filament from ETEC strain 31-10. The densities of water molecules are coloured in red. A part of the map corresponding to one pilin subunit is shown in deep teal. **c**, Ribbon representation of the FimA filament model composing 21 pilin subunits. **d**, Subunit interaction between subunits S (coloured in deep teal) and S_+1_ (coloured in light green). The N-terminal donor strand of one subunit (S is inserted into the hydrophobic groove of the neighbouring subunit (S_+1_) by the so-called “donor-strand exchange” mechanism). **e**, Subunit interaction between subunits S and S_+3_, coloured in magenta and white, respectively. The residues involved in the interactions are labelled and shown as stick models. The water molecules involved in the interaction networks are shown as red spheres. **f**, Molecular packing around subunit S. Each FimA subunit (S) has contacts with a total of six FimA subunits (S_±1_, S_±2_, and S_±3_).

Previous studies have focused on the plasmid-encoded CFs as possible surface-expressed bacterial filaments of ETEC. However, a wide variety of ETEC strains can express chromosome-encoded T1P as an important factor for intestinal colonisation (Sheikh et al., 2017). Since the existence of *fimA* and its associated genes had not yet been examined for ETEC strain 31-10, we determined the whole genome of the strain and identified T1P-related genes (Fig. 5). The FimA sequence was then used for subsequent modelling that resulted in a full-length model of FimA, except for the N-terminal disordered Thr1, confirming further that the observed filament expressed in ETEC strain 31-10 is T1P (Fig. 4c).

**Fig. 5.**
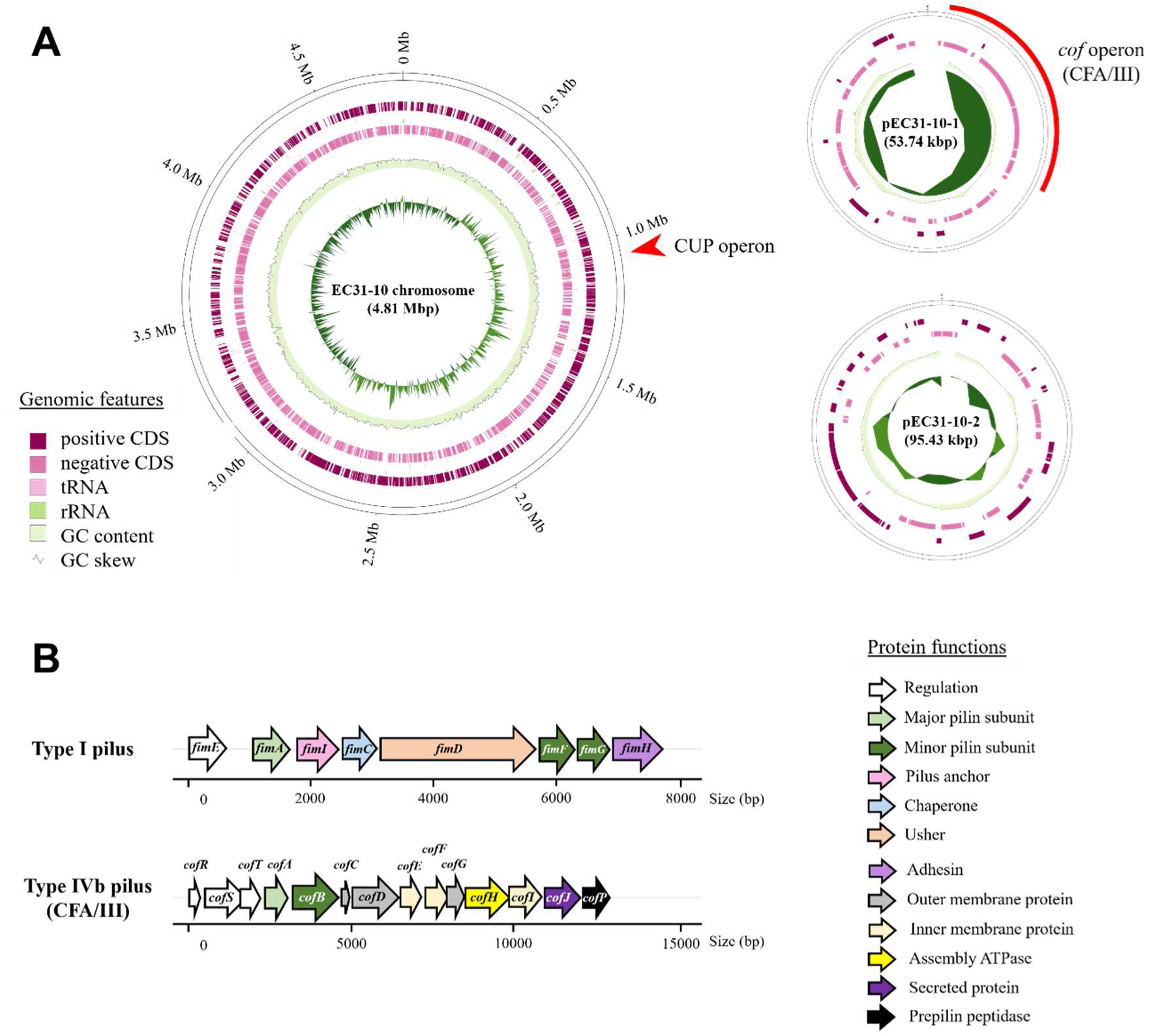
Whole genome sequencing of *E. coli* strain 31-10. a, The genome of enterotoxigenic *E. coli* strain 31-10 visualised using GenoVi (Cumsille, A. et al., 2023) (not to scale). The red arrow and red arcs indicate the position of operons involved in the pili formation of type I and type IVb pilus (CFA/III). b, Gene-cluster of the type I pilus and type IVb pilus (CFA/III) (left panel). Each gene is colour-coded based on the predicted function of the protein. Predicted protein functions for each gene product are shown at the right panel.

The structure of FimA filament is essentially the same as the previously determined cryo-EM structure of T1P from uropathogenic *Escherichia coli* (UPEC), with both taking a right-handed superhelical assembly comprising 3.13 subunits per turn with an axial rise of 8.0 Å per subunit (Hospenthal et al., 2017). Each FimA subunit in the filament adopts an incomplete Ig-like fold, which lacks the C-terminal β-strand that is complemented by the N-terminal donor β-strand of the adjacent subunit (Fig. 4d) (Hospenthal et al., 2017; Waksman and Hultgren, 2009). In addition to this complemented interaction between subunits (S/S_±1_) involving mostly hydrophobic interactions, each subunit (S) primarily forms additional contacts with a total of four subunits, S/S_±2_, and S_±3_, to form a rod-like pilus filament. As expected from their high sequence similarity (91% identity) between the FimA subunit of ETEC and UPEC, almost all of the residues involved in subunit–subunit interactions were conserved. The variable residues were principally observed on the exterior surface of the FimA subunit, which is indicative of immune pressures selecting for antigenic diversification (Appendix Figure. S4) (Spaulding et al., 2018).

The highest-resolution structure for T1P shows that the largest subunit–subunit interactions at the S/S_±3_ interface involving many charged residues located at each of the complementary charged patches, which is thought to be important for pilus assembly (Fig. 4e and Appendix Figure. S5) (Doran et al., 2023; Hospenthal et al., 2017; Spaulding et al., 2018). However, although they are each located close, most of these charged residues do not form salt bridges. These residues include Asp64 and Asp116 (corresponding to Asp62 and Asp114, respectively, in FimA from UPEC), which are proposed to be important in pilus stability (Spaulding et al., 2018). As in the case of the CFA/III filament, the subunit–subunit interface is filled with an extensive number of water molecules, stabilised by the water-mediated hydrogen-bonding networks (Fig. 4f).

## Discussion

Recent advances in cryo-EM and cryo-electron tomography provide powerful tools for identifying and determining the structure of bacterial filaments expressed in response to various types of environmental cues. As demonstrated in this study, cryo-EM is now capable of determining the filament structures at the atomic level, revealing previously uncharacterised pilus biology, including assembly, stability, and flexibility, which are important for efficient infection of pathogenetic bacteria.

Although pilus assembly mechanisms have been well characterised for T1P (Hospenthal et al., 2017; Waksman and Hultgren, 2009), those for T4P have not yet been fully elucidated (Pelicic, 2023). This is partly attributed to the partial melting of α1 observed in almost all of the reported T4P structures that limits the resolution of structural analysis and also prevents the elucidation of the subunit docking process at the inner membrane, where α1 forms a continuous helix. The CFA/III structure with a continuous α1 thus may infer the mechanism of such a process. The structure suggests that each pilin subunit is added to the base of the growing pilus by simple docking (Fig. 6a, ii and iii). The concave binding surface, composed of subunits S_+1_, S_+3_, S_+4_, and S_+7_, of the growing pilus well accommodates the incoming subunit, S, without forcing any conformational changes, even if it has a large globular domain like CofA. Docking is primarily driven by the long-range electrostatic interactions between each pair of charged residues located in oppositely charged surface patches (Figs. 2g and 6b).

**Fig. 6.**
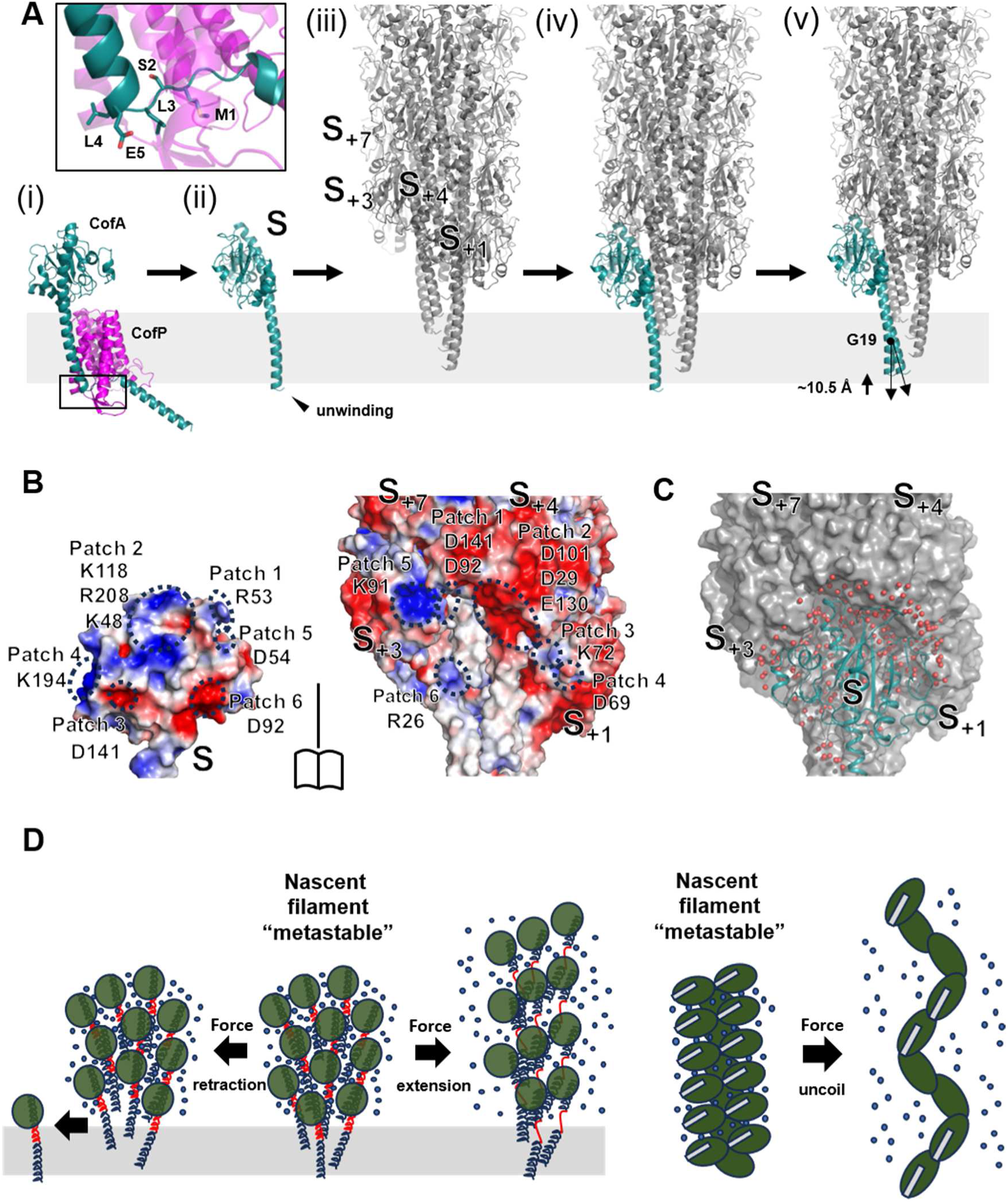
Proposed mechanism of subunit docking in the growing T4bP filament. **a**, A schematic representation of the initial subunit docking process at the inner membrane. After the major pilin subunit is recognised by prepilin peptidase with its N-terminal region (residues 1-5 of matured pilin) partially unwound (i), the type III secretion signal is cleaved off, and the N-terminal five residues may remain unwound and take a turn-like conformation, in which positively charged N-terminal amine is neutralised by the interaction with negatively charged Glu5 (ii). The subunit is then docked onto the base of the growing pilus filament driven by long-range electrostatic interactions among their globular domains (iii and iv). Subsequently, the subunit is thrust up by ∼10.5Å via the function of a cytoplasmic assembly ATPase (v). As described in the main text, the neutralisation of the N-terminus lowers the energetic barrier of this extraction step of the major pilin, with the help of platform proteins. Once extracted, the α1 bends slightly to optimise the hydrophobic and hydrogen-bonding interactions among α1-Ns in the filament core, in which conserved Gly19 in T4b pilin (corresponding to Pro22 in T4a pilin) plays a pivotal role (v). **b,** Electrostatic surface complementarity between the incoming subunit (S) and the base of the growing pilus filament. The electrostatic surface potential was calculated by the *Adaptive Poisson-Boltzmann Solver* (APBS) program (Jurrus et al., 2018). Each pair of oppositely charged patches is indicated by a dashed circle labelled from patch 1 to patch 6. The charged residues located at each patch are also labelled. **c,** Water molecules located at the interface between the incoming subunit (S) and the base of the growing pilus filament. The water molecules are shown as red spheres. **d**, Models of pilus conformational changes upon responding to external forces. Nascent T4P is metastable, in which each subunit interacts with each other through weak non-specific electrostatic interactions stabilised by interfacial water molecules, and easily dissociates or extends under the influence of external forces (left panel). **T**he helically arranged T1P coil is uncoiled because of external forces, but the N-terminal mediated tight interaction between subunits connects each subunit and inhibits its dissociation (right panel).

It has been proposed that complementarity between a conserved, negatively charged Glu5 of the incoming subunit and a positively charged N-terminal amine of the terminal subunit in a growing pilus is a possible driving force for the docking (Craig et al., 2019; Kolappan et al., 2016).

However, our high-resolution structure unambiguously reveals that Glu5 interacts with the backbone amide of Leu3 in the neighbouring subunit (S_+1_) and intramolecularly neutralises the N-terminal amine, by which the N-terminal five residues adopt a turn-like conformation. This neutralisation is suited for their transition to the acyl phase of the membrane. However, as they are initially responsible for anchoring α1 to the inner membrane by interacting with lipid head groups, the question arises regarding the formation of this conformation after the signal sequence cleavage. To obtain some insight, we constructed an AlphaFold-modelled structure of a major pilin–prepilin peptidase complex that showed that the corresponding N-terminal region should be unwound to interact with the peptidase active site (Fig. 6a, i and Appendix Figure. S6). We thus speculate that upon cleavage of the signal sequence, the N-terminal region maintains its unwound conformation by forming a turn-like structure and acts as a trigger to transfer the N-terminus of α1 from the cytoplasm to the acyl phase of the membrane with the help of a cytoplasmic assembly ATPase and/or platform proteins (Nivaskumar et al., 2016; Pelicic 2023; Santos-Moreno et al., 2017) (Fig. 6a, iii to v).

As mentioned earlier, surface charge complementarity is important for a long-range attraction of pilus docking, and many residue pairs with opposite charges are found at each subunit interface (Figs. 2g and 6b). However, most of these charged residues do not form salt bridges. Surprisingly, we recognised only a few direct interactions among residues and found that the interface is filled with an extensive number of water-mediated networks (Fig. 2g and 6c). This also applies to T1P, in which the largest subunit–subunit interface, S/S_±3_, was dictated by suboptimal electrostatic and water-mediated interactions (Fig. 4e,f). Although the hydrophobic N-terminal interactions consolidate the filament assembly in both pilus types, these non-specific water-mediated interactions among their globular domains are “metastable” and sensitive to external forces, and could play important roles in pilus dynamics, e.g., an assembly or disassembly (retraction) and an extension or bending of T4P (Biais et al., 2010; Chlebek et al., 2021) and a coil-uncoil reaction of T1P (Doran et al., 2023; Spaulding et al., 2018), where water molecules play a lubricating role (Fig. 6).

In T4P, this suboptimal interaction is partly attributed to a surface mismatch between the hydrophilic surface of α1-C of incoming subunit (S) and the hydrophobic patch composed of α1-Ns of subunits (S_+3_, S_+4_, and S_+7_) at the pilus base (Appendix Figure. S7). This creates gaps (gap1 and 2) among subunits that can be filled with water molecules (Figs. 2 and 3). As seen in the previously reported T4aP cryo-EM structures (Bardiaux et al., 2019; Garnett and Atherton, 2022; Kolappan et al., 2016; Neuhaus et al., 2020; Treuner-Lange et al., 2024; Wang et al., 2017), the gaps may function as a compression space that accommodates the melting portion of the nearest α1 (Fig. 3c), when responding to external forces such as shear forces *in vivo* and blotting forces during cryo-EM grid preparation (Armstrong et al., 2019; Doran et al., 2023). The susceptibility to such forces apparently depends on the strength of the interactions among globular domains at the filament surface (Treuner-Lange et al., 2024). Thus, filaments composed of pilin subunits with a larger globular domain and longer variable regions would tend to be more tolerant against forces than those composed of smaller ones, as demonstrated here in the CFA/III filament.

The local resolution map of CFA/III intriguingly shows that the resolution on the filament surface is higher than or comparable to that in the interior, suggesting the rigidity of the surface exposed globular domains (Appendix Figure. S8). This feature is also observed in T1P, which has a relatively large globular domain among CUP family filaments (Appendix Figure. S9) (Doran et al., 2023). Given that the water molecules at the subunit interfaces are trapped under cryogenic conditions and contribute to freezing the mobility of protein atoms around them, we thus envision that this “solvent slaving” effect (Stachowski et al., 2022), coupled with densely packed pilins, consolidates the filament and leads to the highest-resolution cryo-EM structures reported to date for each of the pilus types. In CFA/III, it should be noted that the resolution of the middle of α1, including Gly11, Gly14, and Gly19, conserved in the T4bP family, is lower than the others, suggesting potential flexibility. Considering the lubricating role of hydrated water molecules, the filament can be flexible at room or higher temperatures, as captured by its negatively stained TEM images (Appendix Figure. S1) (Kawahara et al., 2016; Kolappan et al., 2012; Taniguchi et al., 2001).

In summary, the high-resolution cryo-EM structures of CUP and T4P purified from ETEC strain 31-10 reveal that, in contrast to the stable hydrophobic filament cores, the interactions among globular domains are formed by rather weak electrostatic interactions stabilised by an extensive number of water molecules that form a hydrogen-bonding network, distributed throughout the filament including even at the inner filament core, which play an important role in pilus assembly, stability, and flexibility. These features depend largely on the size, sequence, and structure of the variable regions of the globular pilin domain that have evolved in bacteria to adapt and/or select their host niche location. As bacterial filaments are crucial components for successful host infection, this structural information will be useful for drug discovery and vaccine development against enteric pathogens with multi-drug resistance.

## Methods

### Pili purification

ETEC strain 31-10 (Taniguchi et al., 2001), pre-cultured in LB medium at 20 °C, was applied to a CFA agar plate (1% casamino acids, 0.15% yeast extract, 0.005% MgSO_4_, 0.0005% MnCl_2_, and 2% agar) (Evans et al., 1977) and incubated at 37 °C to induce the expressions of CFs. The cultured bacteria were collected in phosphate-buffered Saline (PBS) containing 5 mM EDTA, and the expressed bacterial filaments were harvested by vortexing. After centrifugation twice at 8,000 *× g* and 4 °C for 30 min, to remove the bacterial debris, ammonium sulphate was added to 30% saturation to the supernatant to precipitate the filaments. The precipitate was dissolved in PBS and dialysed to remove any remaining ammonium sulphate. For further purification of bacterial filaments, such as CFA/III, we performed density gradient centrifugation at 100,000 × *g* and 4 °C for 3 h using 10%–50% (w/v) sucrose. Ultracentrifugation separated an upper layer containing *E. coli* proteins, a middle layer containing bacterial filaments, and a lower layer containing bacterial debris. The middle layer contains a major pilin CofA of CFA/III, as determined through SDS-PAGE analysis of the fraction. The purified filament fraction was then dialysed against buffer (20 mM Tris-HCl, 150 mM NaCl, pH 8.0) and concentrated to approximately 0.3 and 3 mg/ml for TEM and cryo-EM data collection, respectively.

### Negative-staining transmission electron microscopy

Purified filament fraction (2.5 µl) was incubated at 8 °C for 5 min and applied onto the glow-discharged carbon-coated copper grids, and then washed with two droplets of HEPES buffer (40 mM HEPES-NaOH (pH 7.5)) prior to negative staining with 1% uranyl acetate. The prepared mesh was observed using a JEM-1400 (JEOL Ltd.) operated at 120 kV accelerating voltage and recorded using the built-in CCD camera.

### Cryo-EM data collection

A purified filament fraction (2 µL) was applied to a copper grid (R1.2/1.3 300 mesh; Quantifoil) and glow-discharged for 10 s using a JEC-3000FC Auto Fine Coater (JEOL). The grid was blotted for 3 s at 8°C (100% humidity) using a blot force of 10 and then plunge-frozen in liquid ethane using a Vitrobot Mark IV System (Thermo Fisher Scientific). Screening and data collections were conducted using a CRYO ARM^TM^ 200 or 300 electron microscope (JEOL) operating at 200 or 300 kV, equipped with a cold field-emission gun, an in-column Omega energy filter (20 eV slit width), and a K2 or K3 direct electron detector (Gatan), respectively. For screening, we used the customisable automated data acquisition system JADAS (JEOL), whereas cryo-EM movies for data collection were acquired using the automated EM data acquisition program Serial EM (Mastronarde, D. N., 2003) with 5 × 5 beam-image shift patterns (coma *vs.* image shift calibration was performed prior to data acquisition). A total of 6,899 movies were collected using a nominal magnification of ×60,000 at a pixel size of 0.752 Å/pixel. Each movie was divided into 50 frames using a total dose of 53.77 e^-^/Å^2^, with the K3 detector operating in the correlated double-sampling (CDS) mode. All parameters involved in data collection are presented in Appendix Table S1.

### Cryo-EM image processing for CFA/III filament

All data processing of cryo-EM movies obtained in this study was performed using CryoSPARC v4.5.3 (Punjani et al., 2017). The imported movies were first subjected to beam-induced motion correction by using the function “Patch motion correction,” and the contrast transfer function (CTF) parameters were assessed using “Patch CTF estimation”. We manually picked hundreds of non-overlapping filament segments and performed 2D classification. Selected 2D classes were then used as templates to trace the helical filament found in all the micrographs using the program “Filament Tracer”, and a total of 3,011,705 particles were extracted using a box size of 300 Å. Subsequent 2D classification results in a well-resolved 2D class image that shows a feature corresponding to the T4P filament. A total of 2,887,506 particles were selected for further data processing. We initially calculated the helical volume by “Helical Refinement” without the input of any helical symmetry parameter, and then searched for helical parameters by the “Symmetry Search Utility”, with previously estimated values of an axial rise of 8.5 Å and a twist angle of 96.8° from the negative-staining reconstruction of TCP from *V. cholerae* as a reference (Li et al., 2012), revealed that the volume has a helical rise of 8.01 Å and twist angle of 94.63°. The parameter set was used for “Helical Refinement” that led to a map of 2.11 Å resolution. After local and global CTF refinement, each particle was motion-corrected with “Reference Based Motion Correction” and high-quality particles were then selected by 2D and 3D classification. After further local and global CTF refinement, “Helical Refinement” using 1,425,921 particles yielded a volume with a helical rise of 8.01 Å, a twist angle of 94.55°, and a resolution of 1.78 Å based on the gold-standard Fourier shell correlation. The image processing workflow for the CFA/III reconstruction is depicted in Appendix Figure S2.

### Model building and refinement of CFA/III filament

The initial model of the CFA/III filament was automatically built into the cryo-EM density map using Model Angelo (Jamali et al., 2024), initially without using any sequence input and then using the sequence of major pilin CofA (GenBank code: BAB62897.1) (Taniguchi et al., 2001). The model was manually refined using the program Coot (Emsley and Cowtan, 2004) and subjected to a real-space refinement using phenix.real_space_refine implemented in PHENIX (Afonine et al., 2018). During manual model building using Coot, we observed an alternative conformation of Met1 at lower counter level for both original and sharpened map (at 1.5 and 2.0 RMSD, respectively), in addition to one that directs its N-terminal amine toward the side chain of Glu5 to form an electrostatic interaction (Fig. 2d). The residual density shows that the N-terminal amine is located to interact with surrounding hydrophobic residues of neighbouring subunits, suggesting its methylation (Giltner et al., 2012). However, due to its low occupancy, we did not include this conformation in the final model. Water molecules were automatically located with a criterion of density level (more than 1.5 RMSD) in the original map using Coot and then manually checked. The geometries of the CFA/III filament model were verified using MolProbity (Williams et al., 2018). All refinement statistics are listed in Appendix Table S1.

### Cryo-EM image processing for the FimA filament

Since there are a small number of filaments in the cryo-EM micrographs that apparently have morphology different from CFA/III, we also performed reconstruction of such filaments. The particle picking was done by using the program “Filament Tracer” with a filament diameter of 65 Å. The 2D classification of extracted images suggests that the observed filaments have a feature reminiscent of CUP. The selected 2D classes were then used as templates to trace the filament using the “Filament Tracer”. The filament images observed in all of the cryo-EM micrographs were further segmented into 10,097,033 particles using a box size of 300 Å. After several rounds of 2D classification, 48,324 particles were selected and used in generating an initial helical volume. The search for helical parameters using the “Symmetry Search Utility”, with previously estimated values of an axial rise of 8.0 Å and a twist angle of 115° for type I pili as a reference, revealed that the helical volume has helical parameters with a helical rise of 7.72 Å and twist angle of 114.90°. After local and global CTF refinement and motion correction with “Reference Based Motion Correction”, high-quality particles were selected by 2D classification. After further local and global CTF refinement, the helical refinement using 43,184 particles yielded a well-resolved volume map amenable to the model of T1P FimA as detailed below with a helical rise of 7.71 Å, twist angle of 114.95°, and resolution of 2.20 Å, based on the Fourier shell correlation. A processing workflow for CFA/III reconstruction is depicted in Appendix Figure S3.

### Model building and refinement of the FimA filament

The initial model was automatically built into the cryo-EM density map using Model Angelo (Jamali et al., 2024), without using any sequence input. The modelling estimates the subunit sequence, which has 84% sequence identity with the sequences of the type I fimbrial subunit protein FimA from *Shigella flexneri* and ETEC (GenBank code: UMV06456.1 and QED7455.1, respectively) from UPEC. Since no FimA sequence for ETEC strain 31-10 had been reported, we determined the sequence here (as described below) and then used it for further model building by Model Angelo. The model was manually refined using the program Coot (Emsley and Cowtan, 2004) and then subjected to a real-space refinement using “phenix.real_space_refine” implemented in PHENIX (Afonine et al., 2018). Water molecules were automatically located with criteria of density level (more than 1.5 RMSD) in the original map using the program Coot (Emsley and Cowtan, 2004), and then manually checked. The geometries of the CFA/III and FimA models were verified using MolProbity (Williams et al., 2018). All refinement statistics are listed in Appendix Table S1.

### Genome sequencing and assembly of ETEC strain 31-10

The ETEC strain 31-10 was cultured in LB medium at 25 °C, and cell pellets were collected by centrifugation. Bacterial DNA was extracted from the precipitation using the DNeasy PowerSoil Kit (QIAGEN). The extracted bacterial genomic DNA was sequenced using both short and long-read sequencing platforms. Short read sequencing by MiSeq (Illumina) was performed using the KAPA Hyper Plus Library Preparation Kit (Kapa Biosystems) in a 2 × 151 bp paired-end run. Long-read sequencing data was obtained using MinION (Oxford Nanopore Technologies) sequencing with the Ligation Sequencing Kit SQK-LSK109 (Oxford Nanopore Technologies). Long-read data were assembled using Flye software v2.9.4 (Kolmogorov et al., 2019), resulting in three contigs corresponding to one chromosome and two plasmids. The assembled data were corrected using medaka software v1.12.0 (https://github.com/nanoporetech/medaka) and then minimap2 v2.28 (Li, 2018) and pilon v1.24 (Walker et al., 2014) utilising short-read sequencing data. We annotated the genome of ETEC strain 31-10 using DFAST (Tanizawa et al., 2018).

## Acknowledgements

We thank Hiroki Tanino, Tomoko Miyata, and Keiichi Namba for assistance with cryo-EM data screening at Osaka University. Conventional TEM observations based on negative staining were performed in the Analysis and Development System of Advanced Materials (ADAM) collaborative research facility at the Research Institute for Sustainable Humanosphere, Kyoto University. This research was partially supported by the Research Support Project for Life Science and Drug Discovery (Basis for Supporting Innovative Drug Discovery and Life Science Research (BINDS)) from AMED under Grant Number JP22ama121003. The cryo-EM experiments for data collection were performed with the approval of the SPring-8 Proposal Review Committee (2021B2538, 2022A2538, 2022A2747, 2022B2747, 2023A2738, and 2023B2738). This work was supported by JSPS KAKENHI grant numbers 21K07024 and K10218 (to K.K.), 20K16245 and 23K14519 (to H.O.), 24KJ1623 (to M.I.). This work was conducted as part of “The Nippon Foundation – Osaka University Project for Infectious Disease Prevention”. This work was also supported by AMED under grant number JP223fa627002 (to S.N.).

## Author contributions

**Kazuki Kawahara:** Resources; Data curation; Formal analysis; Validation; Investigation; Visualization; Methodology; Writing—original draft; Writing—review and editing. **Hiroya Oki:** Resources; Data curation; Formal analysis; Validation; Investigation; Visualization; Methodology; Writing—original draft; Writing—review and editing. **Minato Iimori**: Resources; Data curation; Formal analysis; Validation; Investigation; Visualization; Methodology. **Ryuki Muramoto**: Resources; Data curation; Formal analysis; Validation; Investigation; Visualization; Methodology. **Tomoya Imai:** Data curation; Formal analysis; Validation; Investigation; Visualization; Methodology; Writing—review and editing. **Christoph Gerle:** Data curation; Formal analysis; Validation; Investigation; Visualization; Methodology; Writing—review and editing. **Hideki Shigematsu**: Data curation; Formal analysis; Validation; Investigation; Visualization; Methodology; Writing—review and editing. **Shigeaki Matsuda**: Data curation; Formal analysis; Validation; Investigation; Visualization; Methodology; Writing—review and editing. **Tetsuya Iida**: Data curation; Formal analysis; Validation; Investigation; Visualization; Methodology; Writing—review and editing. **Shota Nakamura:** Resources; Data curation; Formal analysis; Validation; Investigation; Visualization; Methodology; Writing—original draft; Writing—review and editing.

## Data availability

The cryo-EM map and model of CFA/III have been deposited to EMDB and PDB, with codes EMD-60902 and 9IUF, respectively. The cryo-EM map and model of type I pilus have been deposited to EMDB and PDB, with codes EMD-60903 and 9IUG, respectively. The sequencing data of ETEC strain 31-10 have been deposited to DDBJ with an accession number of PRJDB18453.

